# Temperature dependence of mosquitoes: comparing mechanistic and machine learning approaches

**DOI:** 10.1101/2023.12.04.569955

**Authors:** Tejas S. Athni, Marissa L. Childs, Caroline K. Glidden, Erin A. Mordecai

## Abstract

Mosquito vectors of pathogens (e.g., *Aedes*, *Anopheles*, and *Culex* spp. which transmit dengue, Zika, chikungunya, West Nile, malaria, and others) are of increasing concern for global public health. These vectors are geographically shifting under climate and other anthropogenic changes. As small-bodied ectotherms, mosquitoes are strongly affected by temperature, which causes unimodal responses in mosquito life history traits (e.g., biting rate, adult mortality rate, mosquito development rate, and probability of egg-to-adult survival) that exhibit upper and lower thermal limits and intermediate thermal optima in laboratory studies. However, it remains unknown how mosquito thermal responses measured in laboratory experiments relate to the realized thermal responses of mosquitoes in the field. To address this gap, we leverage thousands of global mosquito occurrences and geospatial satellite data at high spatial resolution to construct machine-learning based species distribution models, from which vector thermal responses are estimated. We apply methods to restrict models to the relevant mosquito activity season and to conduct ecologically-plausible spatial background sampling centered around ecoregions for comparison to mosquito occurrence records. We found that thermal minima estimated from laboratory studies were highly correlated with those from the species distributions (r = 0.90). The thermal optima were less strongly correlated (r = 0.69). For most species, we did not detect thermal maxima from their observed distributions so were unable to compare to laboratory-based estimates. The results suggest that laboratory studies have the potential to be highly transportable to predicting lower thermal limits and thermal optima of mosquitoes in the field. At the same time, lab-based models likely capture physiological limits on mosquito persistence at high temperatures that are not apparent from field-based observational studies but may critically determine mosquito responses to climate warming.

## INTRODUCTION

Mosquito-borne diseases (e.g., malaria, dengue, Zika, chikungunya, West Nile, yellow fever) are responsible for a significant worldwide burden of infectious disease and represent a major threat to global public health (Bhatt et al., 2013; Gaythorpe et al., 2021; Murray et al., 2012; Puntasecca et al., 2021; Stanaway et al., 2016). As small-bodied ectotherms, mosquito vectors are sensitive to environmental conditions and, in particular, to abiotic factors such as temperature (Mordecai et al., 2019; Paaijmans et al., 2013). Climate change is likely to alter the climatic and habitat suitability for and the geographic distribution of mosquitoes, in turn affecting the distribution of pathogens they transmit (Kraemer et al., 2019; Messina et al., 2019; Parham & Michael, 2010; Rochlin et al., 2013). Understanding the limitations that temperature and other ecological conditions place on mosquito vectors is critically important for predicting how mosquitoes and vector-borne diseases will respond to climate change.

Previous laboratory experiments have measured the relationship between temperature and mosquito life history traits such as biting rate, adult mortality rate, mosquito development rate, and probability of egg-to-adult survival for many vector species (Focks et al., 1993; Rueda et al., 1990; Shapiro et al., 2017; Tun-Lin et al., 2000). Together, these trait thermal performance relationships can provide a mechanistic estimate of equilibrium mosquito abundance as a function of temperature (Mordecai et al., 2013, 2019; Parham & Michael, 2010). For mosquitoes for which these relationships have been estimated, the thermal response curves of individual traits follow predictable patterns: they decline to zero at lower thermal minima and upper thermal maxima and peak at intermediate thermal optima, as expected from first principles of physiology and enzyme kinetics (Amarasekare & Savage, 2012; Huey & Berrigan, 2001; Mordecai et al., 2019). In aggregate, modeled population-level mosquito abundance has a similar unimodal relationship with temperature (Mordecai et al., 2013), reflecting the underlying life history traits (Parham & Michael, 2010). These studies confirm a core tenet of the metabolic theory of ecology, which states that organismal physiology operates within and is restricted by thermal limits (Brown et al., 2004). Temperature-dependent effects of these life history traits in turn affect transmission of mosquito-borne disease (Craig et al., 1999; Johansson et al., 2014; Kushmaro et al., 2015; Lafferty, 2009; Liu-Helmersson et al., 2014; Mordecai et al., 2013, 2017; Morin et al., 2015; Paull et al., 2017; Shocket et al., 2018, 2020; Tesla et al., 2018; Villena et al., 2022; Vogels et al., 2017; Wesolowski et al., 2015). Despite these clear predictions from ectotherm physiology and laboratory thermal performance experiments, it remains unknown to what extent these experimental, laboratory-based mosquito thermal responses predict the realized thermal responses of mosquito populations in the field. In particular: (1) is temperature an important predictor of mosquito distributions? If so, (2) are effects of temperature on mosquito occurrence nonlinear? and if so, (3) do the predicted thermal optima and limits from laboratory experiments align with thermal optima and limits measured in the field?

The rapid rise of biodiversity data, machine learning, and open source satellite imagery in the past decade has enabled new types of approaches necessary to address these questions. Species distribution models offer a way to connect data on species occurrences with environmental covariates to understand how environmental conditions influence the probability of a species occurring in a given location (i.e., habitat suitability). Due to the challenges of accurately detecting a species’ absence, species distribution modeling typically compares locations in which a species has previously been identified (occurrences) and an artificially-generated set of background points (pseudo-absences) which approximate the potentially-habitable and accessible area for a given species (Barve et al., 2011; Elith & Leathwick, 2009; Peterson & Soberón, 2012; Soberon & Peterson, 2005). Species distribution models have been used to characterize the ecological niches (i.e., habitat and climatic requirements) of organisms across the tree of life, including elephants in South Asia (Kanagaraj et al., 2019), lynxes in Canada (Peers et al., 2012), ants in Australia (Leahy et al., 2020), and rare plants in California (Gogol-Prokurat, 2011). Beyond predicting geographic ranges, these models can indicate which environmental covariates are most important for predicting species occurrences, and the functional relationships between environmental conditions and species distributions.

Species distribution models often rely on remotely sensed satellite imagery to provide information about biotic, abiotic, and anthropogenic variables. Recent advances in storage and processing of remotely-sensed imagery, including publicly-available platforms like Google Earth Engine (Gorelick et al., 2017), have increased the resolution and spatial extent over which these models can feasibly be run. In parallel, the Global Biodiversity Information Facility (GBIF) leverages crowdsourced data collection and digitized biodiversity data, where occurrence points of many species are compiled within a central repository (Edwards, 2004). Together, these innovations provide ripe new avenues and abundant data through which to tackle ecological questions.

We focus on the vectors of the world’s highest-burden mosquito-borne diseases (malaria, dengue, chikungunya, Zika, West Nile, and other arboviruses)—*Aedes aegypti, Ae. albopictus, Anopheles gambiae, An. stephensi, Culex pipiens, Cx. quinquefasciatus,* and *Cx. tarsalis*—each of which has well-characterized thermal performances curves in laboratory settings and a wealth of data on occurrences in the field (Kraemer et al., 2015; Messina et al., 2015, 2019; Mordecai et al., 2013, 2017, 2019; Shocket et al., 2018). *Ae. aegypti* and *Ae. albopictus* are important vectors of dengue, chikungunya, yellow fever, and Zika viruses. *An. gambiae* and *An. stephensi* primarily vector the *Plasmodium* spp. protozoans that cause malaria. *Cx. pipiens, Cx. quinquefasciatus,* and *Cx. tarsalis* transmit West Nile, St. Louis encephalitis, Western Equine encephalitis, and other zoonotic viruses.

While previous mosquito species distribution models have focused on estimating habitat suitability and predicting occurrence (Kraemer et al., 2019; Messina et al., 2015, 2019), our goal is different: to specifically dissect the relationship between temperature and probability of mosquito occurrence, while incorporating other drivers of occurrence, and to compare this relationship among mosquito species and with laboratory-based model predictions. For this reason we created new models, rather than using published ones, that use consistent methods across multiple mosquito species. Further, we aimed to construct species distribution models for each focal species using spatially- and temporally-similar data sources and consistent model assumptions in order to compare thermal dependence across species.

Toward these objectives, this project leverages new computational and methodological advances to address a long-standing question in disease ecology: does temperature predict realized mosquito distributions in the field, and if so, what are the functional relationships? We aim to compare the insights from a ‘top-down’ machine learning and statistical modeling approach to results from ‘bottom-up’ models of mosquito population sizes from trait-based experiments in the lab (Mordecai et al., 2019) to discern whether and how results from the lab setting can translate into the real world. We make two important methodological advances to ensure that our species distribution models capture true thermal limits on mosquito occurrence, rather than sampling bias or restricted geographic ranges that covary with temperature. First, we sample background pseudo-absence points only from the set of ecoregions in which the focal mosquitoes occurred, as well as adjacent ecoregions. This ensures that comparator pseudo-absence points are selected from regions where the focal mosquito species could have realistically been found but were not, or in other words, zones that are ecologically plausible for an occurrence. Second, we restrict temperature measurements to an ‘activity season’ during which each mosquito species is blood-feeding and reproducing, and not in dormancy or torpor. This captures temperature constraints within a physiologically realistic period, rather than constraints due to overwintering or drought persistence, which are less likely to be comparable to laboratory-based trait thermal response experiments. By using global mosquito occurrences from 2000-2019, geospatial satellite-derived covariates at high spatial and temporal resolution, bias-reducing methods described above, and a gradient-boosted classification tree machine learning algorithm well-suited for prediction tasks, we aim to provide evidence that spans across continents and decades for the globally important vectors of human malaria and arboviral disease. With a greater understanding of thermal constraints on vector species occurrence, we can improve the ability to accurately project and mitigate the future impacts of climate change on mosquito-borne disease distributions.

## METHODS

### Overview and study period

We focused on 2000-2019 to isolate recent land-use and climate patterns but provide a long enough time period to capture mosquitoes’ stable spatial distribution. The environmental covariates and mosquito occurrences were both extracted for this time period, providing temporal consistency in the analysis (Wisz et al., 2013). An overview of the methods can be seen below (Fig. 1). Computation was performed in R v4.1.1, Google Earth Engine and the Sherlock computing cluster at Stanford University.

**Figure 1.**
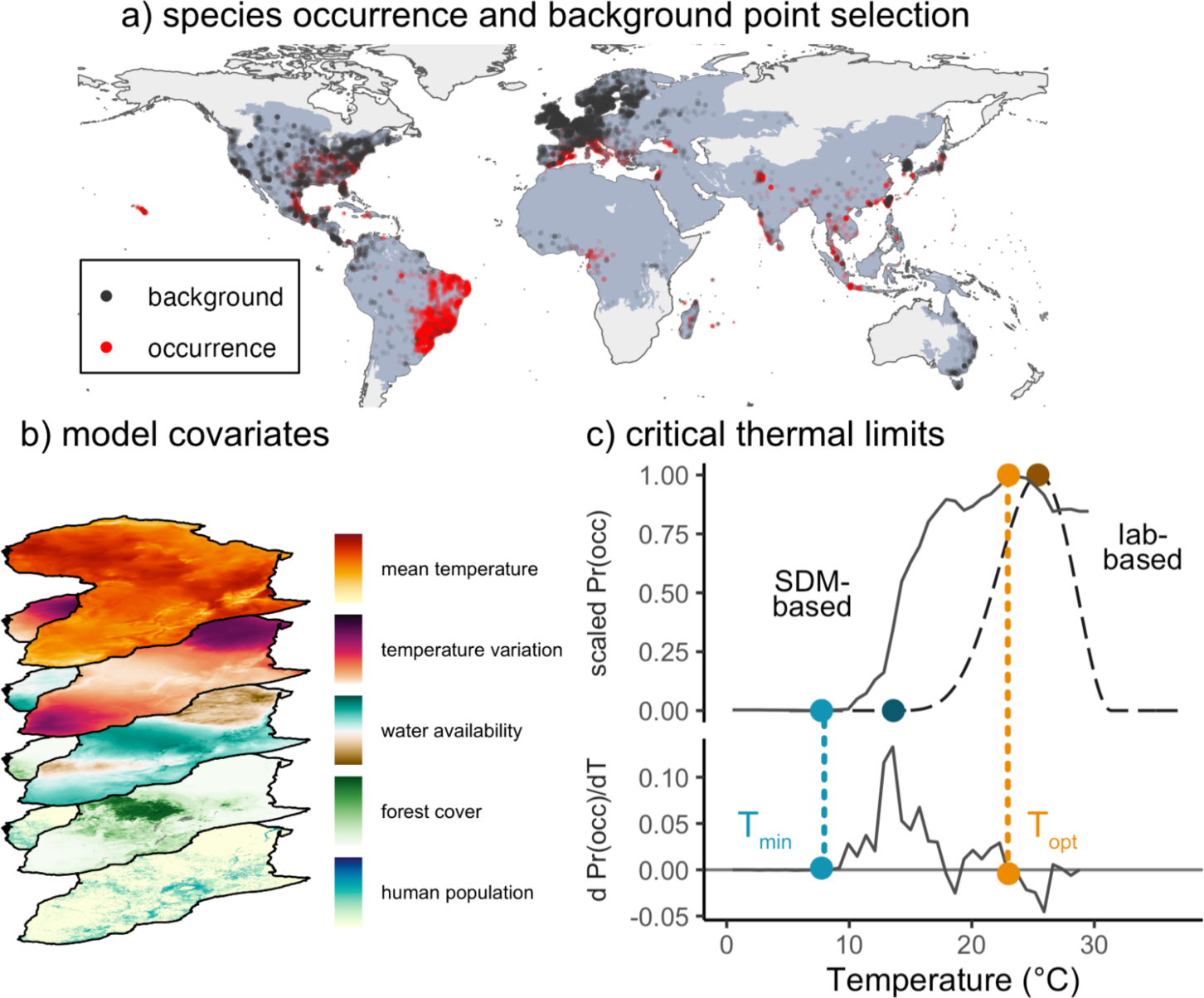
Methods for comparing mosquito thermal performance derived from species distribution models (a-c) and laboratory trait-based models (c) The analysis involved two major steps: statistical modeling of mosquito occurrence using global species occurrences (a) and geospatial data (b) to identify the relationship between temperature and species occurrence (solid line, c), and comparison with mosquito abundance curves previously derived from laboratory life history traits (Mordecai et al., 2019) (dashed line, c). Temperature-dependent mosquito abundance (M(T); dashed line in c) is modeled as a function of temperature-dependent eggs laid per female mosquito per day (EFD(T)), probability of egg-to-adult survival (pEA(T)), development rate from egg to adult (MDR(T) = 1/development time in days), and daily adult mosquito mortality rate (***μ***(T)). Trait curves adapted from Mordecai et al., 2019 and Villena et al., 2022; M(T) equation originally derived from Mordecai et al., 2013. Thermal minima and optima of the probability of occurrence from the species distribution models were identified by calculating the temperature at which the empirical derivative was first positive and stayed positive for the next step (light blue point, c) and the temperature where the empirical derivative was zero and the probability of occurrence was at its maximum (light orange, c). Sources of data for the stack of covariates are described in Table S2.

### Environmental predictors

Environmental covariates were defined *a priori* in order to avoid data-dredging (Rochlin et al., 2013), and were based on a literature search of previous species distribution models. Variables were informed by ecological and biological relevance, expectation to play a role in mosquito vector ecology, and top contributor status in previous mosquito species distribution models (Akpan et al., 2018; Conley et al., 2014; Fatima et al., 2016; Mweya et al., 2013). Environmental covariates were computed using remotely sensed and reanalysis data, and resampled using bilinear interpolation to consistent 1 km x 1 km resolution on Google Earth Engine. This approach using global satellite data avoids the spatial interpolation limitations of common climatic datasets such as WorldClim, in which data are sourced from ground meteorological stations with uneven geographic distribution and nonrepresentative selection.

In order to verify minimal overlap between predictor variables, a pairwise correlation analysis was performed with a Pearson’s correlation coefficient r < |0.8| threshold (Fig. S1). Annual averages for covariates were computed across the study period, providing the characteristic climatic conditions across the study’s temporal range.

### Temperature predictors

While many species are active year-round (Table 1), numerous species physiologically undergo photoperiodic diapause in the winter months when ambient light is low in order to conserve energy for life sustenance (Zhang et al., 2019) or aestivate during the dry season when no breeding habitat is available (Lehmann et al., 2010). To capture the temperature that mosquitoes experience while active, temperature covariates were calculated over the activity season.

**Table 1.** Mosquito species by activity season.

| Mosquito Species | Activity Season (Reference) | Raw Number of Occurrences | Unique Cell Centroids |
| --- | --- | --- | --- |
| *Aedes aegypti* | Year Round (Chang et al., 2007; Mitchell, 1995) | 37,115 | 9,299 |
| *Aedes albopictus* <sup>#</sup> | Photoperiod (Lacour et al., 2015; Poelchau et al., 2013) | 39,488 | 8,688 |
| *Anopheles gambiae* | Precipitation (Lehmann et al., 2010; Yaro et al., 2012) | 13,650 | 903 |
| *Anopheles stephensi* | Year Round (Sinka et al., 2011) | 1,232 | 358 |
| *Culex pipiens* | Photoperiod (Rinehart et al., 2006; Robich & Denlinger, 2005) | 98,276 | 1,949 |
| *Culex quinquefasciatus* | Year Round (Almirón & Brewer, 1996; Meuti et al., 2015) | 30,978 | 2,670 |
| *Culex tarsalis* | Photoperiod (Buth et al., 1990) | 44,496 | 909 |

<sup>#</sup> This species enters winter diapause in some regions such as the Northeast US, but not other regions like the Southeast US.

For the purposes of this study, the photoperiodic activity season was defined as the time period encompassing days with at least 9 hours of sunlight (Fig. S2). This definition of the activity season was chosen based on a variable separate from temperature and on laboratory studies that found that either 8 or 9 hours of light can induce diapause in mosquitoes in the absence of temperature change (Poelchau et al., 2013; Rinehart et al., 2006; Robich & Denlinger, 2005). Other tropical mosquito species (e.g., *Anopheles gambiae*) do not enter photoperiodic dormancy due to their equatorial setting. Rather, these species’ activity seasons are instead constrained by lack of precipitation, when insufficient habitat is available for immature development and survival. Studies have highlighted how Sahelian *Anopheles gambiae* mosquitoes virtually disappear in the dry season then re-establish in the wet season through aestivation behavior (Lehmann et al., 2010; Yaro et al., 2012), and climatological studies in the Democratic Republic of the Congo characterize the dry season as having less than 50 mm of rain in the past month (Kazadi & Kaoru, 1996). Precipitation activity season is therefore defined in our study as the time period in which there is, on average, at least 50 mm of precipitation in the past 30 days (Fig. S3).

Predictors capturing thermal central tendency include year round temperature mean, photoperiod activity season temperature mean, and precipitation activity season temperature mean. Measures of seasonal thermal variation include year round temperature standard deviation, photoperiod activity season temperature standard deviation, and precipitation activity season temperature standard deviation.

### Other environmental predictors

Environmental variables other than temperature included measures of vegetation cover, such as enhanced vegetation index mean, enhanced vegetation index standard deviation, and forest cover percentage, to capture land cover that might affect mosquito habitat suitability (Table S2, Fig. S4). Human population density and cattle density were included to capture sources of blood meal, anthropophilic hotspots, and human-seeking behavior. Precipitation of the driest quarter, precipitation of the wettest quarter, and surface water seasonality were included to capture hydrology-driven characteristics influencing mosquito breeding site availability or abundance, while wind speed was included to capture potential aerial dispersal limitation of mosquitoes. Precipitation of the driest quarter and precipitation of the wettest quarter were defined as the set of 3 consecutive months with the least and most precipitation, respectively, across the 20-year study period.

### Occurrence and background

#### Presence occurrences

Focal mosquito species were selected based on global disease burden, availability of life history trait data from published laboratory studies, and abundance of publicly-accessible species occurrences, resulting in seven species: *Ae. aegypti, Ae. albopictus, An. gambiae, An. stephensi, Cx. pipiens, Cx. quinquefasciatus,* and *Cx. tarsalis*. Collectively, these seven species span nearly all habitable continents (Fig. S5-S11). Species occurrence records for each species were obtained from the Global Biodiversity Information Facility (GBIF) for the years 2000-2019. Supplemental occurrence points were obtained from two papers for *An. stephensi* and *An. gambiae,* which had low sample sizes from the GBIF database (Sinka et al., 2020; Wiebe et al., 2017). Occurrences explicitly tagged in Africa were discarded from the supplemental *An. stephensi* data, as these are part of an ongoing species invasion outside of the native range of the species in South Asia (Sinka et al., 2020), and species distributions models have limitations in detecting environmental suitability for invasive species as they do not yet occupy their environmental niche and are not yet in equilibrium with their environment (Barbet-Massin et al., 2018).

Raw datasets were cleaned to remove occurrences with unknown basis of record and those obtained through fossil records, due to the temporal and geographic uncertainty between when and where the organism actually existed during its lifetime and its location of fossilization. Uncertainty in the latitude and longitude coordinates of occurrence points were taken into account through a two-step process. First, points with a coordinate uncertainty less than 1000 meters—the diameter of our environmental covariate grid cell size—and those points with missing coordinate uncertainty were selected. Of the points with missing coordinate uncertainty, we retain only those points with a latitude and a longitude with at least two significant decimal points (i.e., each hundredth corresponds to approximately 1.11 km). The occurrence data were then restricted to points that reside on landmasses and not in the ocean, those with non-missing values for all covariates, and, when relevant, and those that occurred in areas where the species’ activity season length was greater than 0 (i.e., when the precipitation was above a certain value in the grid cell, i.e., precipitation activity season, or when there was a certain amount of daylight hours in the grid cell, i.e., photoperiod activity season) (Fig. S2-3). We filtered the occurrences to the cell centroids of the unique 1 km x 1 km cells in which occurrence points were obtained, which serves to spatially thin the occurrence points and reduce the effect of biased sampling (Kramer-Schadt et al., 2013) (Table 1; Table S1).

#### Pseudo-absence background sampling

Species distribution models work by comparing the environmental covariates that best predict locations with species occurrences to those in which the species does not occur, to understand the environmental conditions that constrain a species’ distribution. As the majority of information is on species occurrences rather than species absences, methods for species distribution modeling have developed to sample pseudo-absences: points that represent the area in which a species could have been sampled yet was not reported. In order for these sampled points to serve as plausible points where the species of interest was not found, they should be within the species’ accessible area, or the region that is reasonably reachable by the species through dispersion or migration over the relevant time period (Barve et al., 2011; Peterson & Soberón, 2012; Soberon & Peterson, 2005). Choosing pseudo-absences that occur within the same ecoregions, i.e., geographic regions that possess similar species and community assemblages, as occurrences provides a way to geographically restrict possible background points. Moreover, these points should be chosen in a way that takes into account sampling bias and collection effort.

To sample background points, we first created a bias mask to probability-weight the sampling. 72 million occurrences from Class Insecta, filtered to the same criteria as mosquito occurrence points (i.e., <1000m coordinate uncertainty or at least two significant decimal points of latitude and longitude, 2000-2019 study period, no fossil record or literature points), were extracted from GBIF. In order to capture the frequency of GBIF sampling conducted in a specific geographic location, the number of insects were tallied for each 1 km x 1 km grid cell, and these counts were converted into proportions of all insects. This proportion was regarded as the probability weight for that given grid cell. To provide adequate sample size while avoiding a skew of the overall sample towards disproportionately favoring pseudo-absences over occurrences, we selected a number of unique background cell centroids equal to twice the number of unique occurrence centroids (Liu et al., 2013). Background cells were sampled at random without replacement and weighted by the bias mask from the geographical region that was bounded by the focal species’ ecoregions of occurrence. In addition, we include adjacent ecoregions in the background sampling space to promote contiguity in the potential ecological spaces from which pseudo-absences can be drawn and to better encapsulate the broad range of environmental covariates that a given species may experience in nature. Ecoregions were defined as RESOLVE Ecoregions that intersected with a given species’ occurrence centroids, buffered with a 200 km radius buffer that approximated an upper end of the wind-assisted geographic mobility range of a mosquito (Chevillon et al., 1995; Verdonschot & Besse-Lototskaya, 2014). By buffering occurrences and then choosing the set of ecoregions and adjacent ecoregions in which these buffered points lie, we approximate what is known in ecological theory as the ‘accessible area,’ described above (Barve et al., 2011; Peterson & Soberón, 2012; Soberon & Peterson, 2005). Because our focal questions centered on species’ thermal breadth, we wanted to ensure that as broad a temperature range as was reasonable was included in the background sampling. Specifically, we checked whether our background sampling scheme provided a temperature distribution at least as wide as that of the occurrences—a consideration important to ensure that the statistical model can ascertain thermal minima or maxima.

### Species distribution modeling

#### XGBoost model fitting

Raster values for each occurrence and background centroid were extracted from the stack of covariates for a given species. The data was partitioned into a training and evaluation set, where 80% of the data was randomly selected for the training set and 20% was randomly selected for the validation set. We used stratified sampling so that there was an equal proportion of presence points in each set. We used gradient boosted classification trees to model the training data, as this class of algorithms fits flexible and nonlinear relationships (including thresholds) between environmental conditions and probability of species occurrence, identifies interactions between covariates due to the structure of the trees, and has been successfully used in other species distribution models (Kraemer et al., 2015; Li et al., 2022; Messina et al., 2019). Gradient boosted trees are a type of supervised learning algorithm that iteratively fits trees, each time fitting to the residual errors from the previous predictions, effectively ensembling weaker learners to accurately predict a target variable (Natekin & Knoll, 2013). We fit models using eXtreme Gradient Boosting (XGBoost) (Chen et al., 2015) in R. Among the strengths of the XGBoost algorithm are its rapid computational speed when dealing with large-scale data, ability to handle class-imbalanced training sets (i.e., the number of observations are not equal across each level of the outcome variable), and ability to tolerate highly-collinear predictor variables. To quantify variation around model performance and output, we fit the model with 20 random train and validation splits.

#### Bayesian hyperparameter optimization

Gradient boosted classification trees rely on hyperparameters that control how each tree is fit and how individual trees’ estimates are combined. Bayesian optimization was conducted to tune key hyperparameters initialized with a range of user-inputted values, including learning rate (range: 0.0001 to 0.3), maximum tree depth (range: 2 to 50), minimum child weight (range: 1 to 50), subsample of observations used in each tree (range: 0.25 to 1), subsample of columns to use in each tree (range: 0.5 to 1), and minimum split loss (range: 0 to 100) (Chen et al., 2015; *XGBoost Parameters*, n.d.). For each round of the Bayesian optimization (up to a maximum of 24 rounds), we use 5-fold cross validation where we split the training data into 5 disjoint sets, and train the model on each combination of 4 folds while predicting out-of-sample for the fifth fold. We use the average out-of-sample log loss averaged over all 5 folds to identify the optimal number of boosting rounds (up to 10,000) for each iteration of the Bayesian optimization. After all iterations of the Bayesian optimization, we select the set of hyperparameters with the lowest out-of-sample log loss and train a final model using all 5 folds—representing all of the training data, which is 80% of the full data set.

#### Interpreting the model

Receiver operating characteristic (ROC) curves and their area under the curve (AUC) values, which graphically and numerically illustrate the discrimination ability of a binary classifier, were computed for model predictions on both the training and evaluation sets. We were particularly interested in two aspects of the resulting models: the inferred relationship between the probability of occurrence and temperature, and the relative importance of temperature in predicting a species distribution, compared to other environmental covariates. To understand the relationship between temperature and occurrence, we plotted univariate partial dependence plots for temperature mean and temperature standard deviation. Depending on the species, we use annual temperature, temperature during the photoperiod-dictated activity season, or precipitation-dictated activity season. Partial dependence plots depict the mean value of the probability of occurrence given temperature across models fit to all combinations of other variables at each value of temperature. To assess the importance of temperature in relation to other environmental variables, measured in gain (i.e., the increase in accuracy that a given variable provides when a regression tree branch is split upon that variable) (*XGBoost Parameters*, n.d.), we computed variable importance scores for the full set of predictors and compared them with temperature. We computed variable importance and PDPs for each bootstrapping iterations described above.

### Mosquito abundance curves from the lab

Physiological life history trait thermal performance curves were collected from published literature (Mordecai et al., 2019; Shocket et al., 2020; Villena et al., 2022). Briefly, in previously published work asymmetric responses like mosquito development rate (MDR) were modeled using a Briere function, and both concave-down symmetric responses like eggs per female per day (EFD) and probability of egg-to-adult survival (pEA) as well as concave-up symmetric responses like mortality rate (μ) were modeled using quadratic functions. 5,000 posterior draws were selected from each distribution, and mosquito abundance M(T) was calculated using Equation (1), which was originally derived in Mordecai et al., 2013, adapted from Parham and Michael, 2010. Median and 95% credible intervals were computed for each species’ M(T) and critical thermal limits.

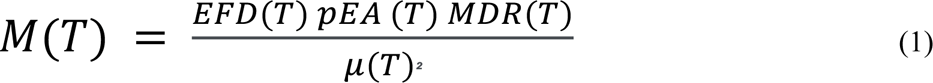

### Comparison of thermal relationships from XGBoost versus lab-based models of mosquito abundance

We compared thermal minima and optima of the partial dependence plots from the XGBoost model to the mosquito abundance M(T) curves from the lab trait trajectories. For species distribution models, thermal minima were identified as the temperature at which the partial dependence plot began increasing, which we operationalized as the first time the empirical derivative was positive and stayed positive for the next step in temperature as well (Fig. 1c). Thermal minima could also correspond to the threshold temperature at which species probability of occurrence is increasing most rapidly, so we supplementarily identify thermal minima as the temperature where the empirical derivative is largest (Fig. S14). Thermal optima were identified as the point where the empirical derivative was zero and the partial dependence plot was at its maximum (Fig. 1c). We did not identify thermal maxima, as few species had partial dependence plots that clearly declined after the thermal optima and then reached a lower plateau that was within the range of observed temperatures. For the mosquito abundance M(T) curves derived from laboratory studies, to determine the thermal minima, we identified the temperature at which M(T) exceeded zero for each M(T) curve calculated from posterior draws and then calculated the median, 2.5th and 97.5th percentiles to produce a central estimate and confidence interval for thermal minima. For thermal optima, we identified the temperature where M(T) was highest for each curve, and similarly calculated median and 95% confidence intervals. We calculate the Pearson correlation between the median species distribution model-based thermal minima and the median lab-based thermal minima, and repeat for thermal optima to quantify how well the two methods compare. In particular, because temperature varies substantially in the field and M(T) curves are based on constant temperature measurements in the lab, we expect nonlinear averaging to affect the exact thermal limits observed in the field.

## RESULTS

The species distribution models predicted the occurrence of all focal species with high accuracy: discrimination across both training (in-sample) and evaluation (out-of-sample) datasets produced an area under the receiver operating curve (AUC) above 0.9, where 1 indicates perfect discrimination of presence and absence and 0.5 is no better than a coin toss (Fig. 2, Fig. S12). Of the focal species, *An. gambiae* recorded the lowest out-of-sample evaluation AUC of 0.938. The fact that the AUCs in the evaluation set were comparable to the training set suggests that the models are learning general patterns, differentiating between training and evaluation sets, and not overfitting to the training data.

**Figure 2.**
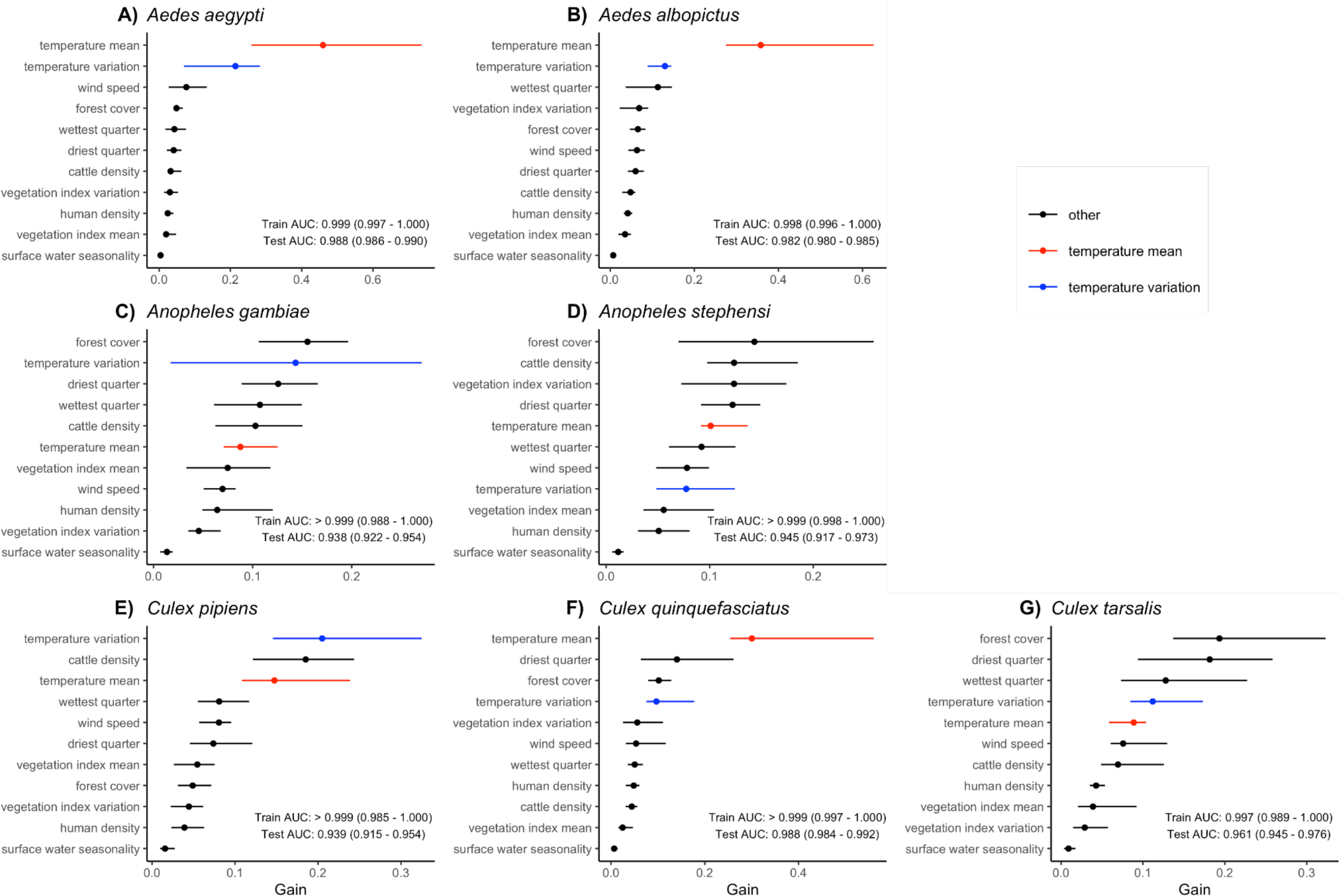
Temperature mean (red) and standard deviation (blue) were important predictors of mosquito occurrence for all focal species. Variable importance, measured in gain, is shown for each predictor variable by species. Temperature mean variables, including year-round temperature annual mean, photoseason activity season temperature mean, and precipitation activity season temperature mean, are colored in red; only the most biologically appropriate temperature mean variable was included for each species (Table 1). Likewise, temperature standard deviation variables (Table 1) are colored in blue. Other predictors (black bars) include cattle density, enhanced vegetation index mean, enhanced vegetation index standard deviation, forest cover percentage, human population density, precipitation of the driest quarter, precipitation of the wettest quarter, surface water seasonality, and wind speed. Points represent the median importance across the 20 bootstrapped model iterations and lines represent the range over all model iterations. Test and training AUC values are similarly medians and full ranges over the model iterations.

Measures of temperature, i.e., either annual or activity season-limited (as appropriate) temperature mean and temperature standard deviation, were consistently important predictors of mosquito occurrence, as measured by gain (Fig. 2). For all species, mean temperature was among the top six predictors. For *Ae. aegypti* and *Ae. albopictus,* temperature mean was the top predictor and temperature standard deviation was second (Fig. 2). *An. gambiae, Cx. pipiens,* and *Cx. tarsalis* had temperature standard deviation as a more important predictor than temperature mean (Fig. 2).

Beyond temperature, different predictors were important for different mosquito species. Forest cover was the most important predictor for *An. stephensi, An. gambiae,* and *Cx. tarsalis*, and precipitation of the driest quarter was the most important predictor for *Cx. quinquefasciatus* (Fig. 2). Precipitation variables were among the top-5 predictor variables for all species. Precipitation of the wettest quarter was among the top-5 predictors for *Ae.aegypti, Ae. albopictus* and *Cx. pipiens* (Fig. 2). Precipitation of the driest quarter was among the top-5 predictors for *An. stephensi* and *Cx. quinquefasciatus*, and both precipitation variables were among the top-5 predictors for *An. gambiae* and *Cx. tarsalis* (Fig. 2) Surface water seasonality, however, was the least important predictor variable for all species distribution models (Fig. 2).

While mechanistic relationships between temperature and mosquito abundance, based on laboratory-derived M(T), were unimodal and hump-shaped in form, the temperature relationships inferred from the statistical models (univariate XGBoost partial dependence plots; PDPs) were typically nonlinear with steep thresholds and plateaus (Fig. 3). Both the mechanistic and statistical models showed steep increases as temperatures exceeded lower thermal limits (Figs. 3). By contrast, while M(T) consistently responded unimodally to temperature, relationships from XGBoost PDPs mostly only showed lower thermal limits and thermal optima, but not upper limits (Fig. 3). The difference in functional forms between M(T) and XGBoost PDPs is consistent with the fact that they each describe distinct processes. The mechanistic model, M(T), describes a relationship between temperature and mosquito *abundance* that continuously varies with temperature and the underlying laboratory-measured traits. By contrast, the PDPs from XGBoost capture the relationship between temperature and mosquito *occurrence probability*, which we expect to rise and fall sharply and reach a plateau at intermediate, suitable temperatures. The M(T) curves for all species showed clear thermal minima, optima, and maxima because the underlying laboratory experiments captured the full range of temperatures at which mosquito performance is optimized and at which it declines to zero.

**Figure 3.**
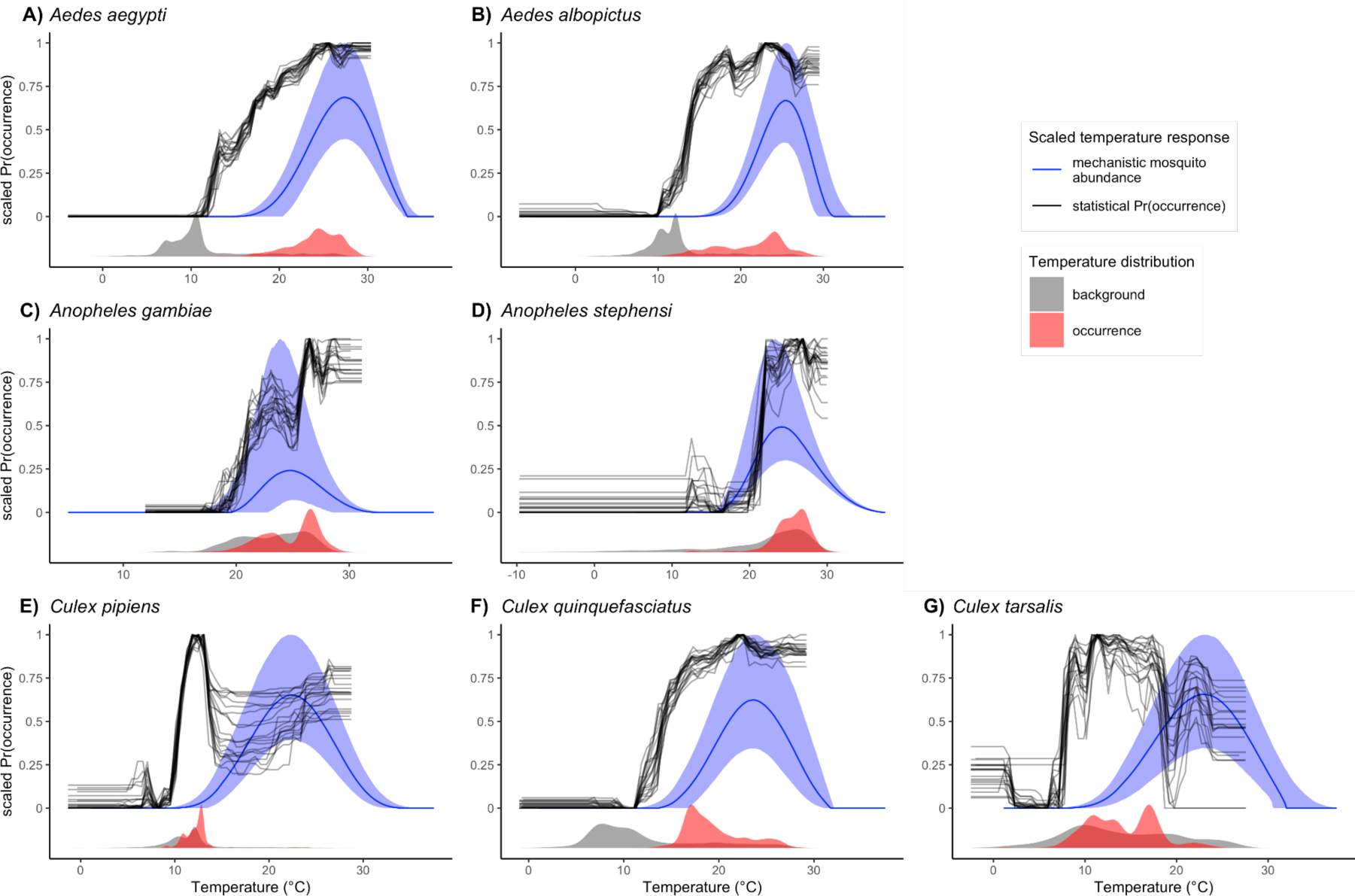
Comparing mosquito temperature responses from mechanistic laboratory-based mosquito abundance models and XGBoost statistical models. Mosquito abundance for each species is estimated as the median M(T) curve with a shaded 95% credible interval band (blue lines and shaded ribbons). XGBoost covariate responses are univariate partial dependence plots (PDPs) for annual mean temperature, where mean temperature was bounded by photoperiod or precipitation for a subset of species, that show the marginal effects of temperature on model prediction (black lines). Each black line for the PDPs represents one of the 20 model iterations. Both M(T) and PDPSs are scaled to range from 0 to 1. Grey and red density plots in the bottom of each panel show the distribution of observed annual temperatures for background and occurrence points, respectively.

Given that both the laboratory-based mechanistic approach (M(T)) and the statistical approach (XGBoost PDPs) predicted thermal minima and optima for each species, we asked whether the results of the two approaches were concordant. The estimates of thermal minima between M(T) and PDP were highly correlated across species (r = 0.902; Fig. 4; Fig. S13, S14), suggesting that M(T) captures key species-specific temperature constraints on occurrence in the field. Mosquito species of the *Aedes* and *Culex* genera exhibited thermal minima that were 2-6°C cooler in the field than predicted in the laboratory, while *Anopheles* mosquitoes exhibited thermal minima 0.5°C higher in the field than in laboratory settings (Fig. 4; Fig. S13, S14). *Cx. tarsalis* and *Cx. pipiens* consistently had the lowest and second-lowest thermal minima, respectively, and *An. gambiae* consistently had the highest thermal minimum when viewed across M(T) and PDP (Fig. 4; Fig. S13-S14). The alternative identification of thermal minima as the threshold temperature with the greatest increase in species occurrence resulted in a similarly high correlation between M(T) and PDP-based thermal minima (r = 0.897), but resulted in thermal minima approximately 5°C warmer in the field than predicted by the laboratory for *Anopheles* mosquitoes (Fig. S14). Thermal optima were similar, although more weakly correlated, between M(T) and PDP estimates (r = 0.685), supporting, at least in part, the conclusion that field based observations capture key components of lab-based non-linear response to temperature. However, the relation appeared to deviate the most for temperature species *Cx. pipiens* and *Cx. taralis,* for which thermal optima were 9.5°C and 10°C lower, respectively, in the field versus laboratory (Fig. 4; Table S3; Fig. S13-S14). We could not compare thermal maxima because our XGBoost models did not identify thermal maxima for any focal species.

**Figure 4.**
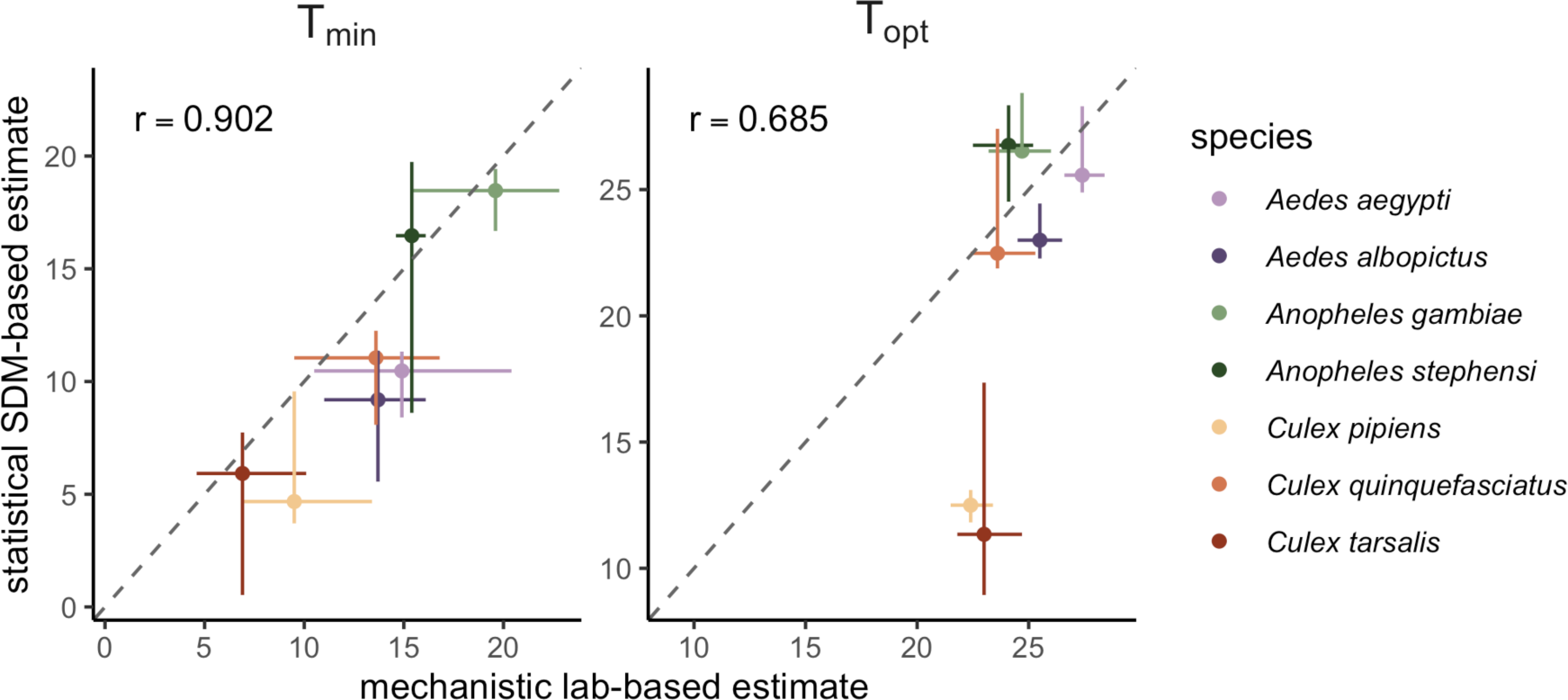
Mechanistic models (M(T); x-axis) captured species’ thermal minima observed in the field (XGBoost PDPs; y-axis) The dashed diagonal line is the 1:1 line, and mosquito species are colored by genus. The Pearson correlation (r) was 0.902 for the PDP versus M(T) relationship for average T_min_ and 0.685 for the PDP versus M(T) relationship for average T_opt_. For mechanistic estimates of T_min_ and T_opt_, points indicate medians and horizontal line ranges are the 95% CI from the 5000 posterior samples used. For statistical estimates of T_min_ and T_opt_, points indicate medians and vertical lines are the full minimum to maximum range over the 20 bootstrapped model iterations.

## DISCUSSION

For all seven major vectors of human disease we investigated, species distribution models captured occurrence probability with high discrimination and accuracy and temperature was an important predictor (Fig. 2, Fig. S12). In particular, temperature mean and standard deviation were highly important predictors across all focal species (Fig. 2). This finding echoes the importance of temperature as a predictor in past models, although many such models define temperature differently from our study (e.g., temperature suitability, temperature of the hottest quarter, temperature of the coldest month) (Brady et al., 2013; Cianci et al., 2015; Fatima et al., 2016; Kraemer et al., 2015). Four species (*Ae. aegypti*, *Ae. albopictus*, *Cx. pipiens,* and *Cx. quinquefasciatus*) had either temperature mean or standard deviation as their most important predictor (Fig. 2). Nonlinear relationships with temperature from ‘bottom-up’ models of mosquito abundance from lab life-history traits were partially recapitulated in the ‘top-down’ statistical models of global-scale occurrence data, particularly at the lower end of the temperature range, consistent with previous work on other mosquito species (Fatima et al., 2016). In general, thermal minima and optima were highly comparable across species between the trait-based and statistical approaches (Fig. 4). However, thermal maxima were not comparable: in particular, XGBoost did not consistently detect thermal maxima for occurrence in the field, despite laboratory evidence that mosquito life history traits decline precipitously at high temperatures.

Thermal physiology theory and experiments have established that organismal performance is constrained to an operative range of temperatures at which key functions can occur and populations can stably persist (Angilletta & Angilletta, 2009; Bernhardt et al., 2018; Dell et al., 2011; Deutsch et al., 2008; Somero, 2002). Yet, realized species distributions may not clearly reflect these thermal constraints if other factors are also important for constraining distributions, including rainfall, seasonality, biotic interactions, habitat, movement rates, and the range of temperatures that occur within accessible regions (Buckley et al., 2010; Crozier & Dwyer, 2006; Davis et al., 1998; Guisan & Thuiller, 2005; Heikkinen et al., 2007; Lounibos & Juliano, 2018; Pulliam, 2000; Wisz et al., 2013). For example, aridity may constrain mosquito distributions beyond direct high-temperature constraints. Likewise, average temperatures across activity seasons may not reach levels high enough to exclude mosquito persistence (Bernhardt et al., 2018; Lambrechts et al., 2011; Paaijmans et al., 2009, 2010, 2013). Alternatively, more extreme temperature variation at low or high means can limit organismal performance, and even moderate variation can reduce performance near optimal temperatures (i.e., where M(T) curves are concave-down; Fig. 3). The impact of temperature variation could explain the observation that for species with cooler thermal minima and optima, estimates at constant temperatures in the lab were up to 10°C warmer than estimates in the field (Fig. 4).

This highlights the importance of climate change projections that consider both field-based estimates of thermal and other constraints on species distributions and more mechanistic estimates, particularly for thermal optima and maxima, which may be difficult to observe in the field under current and historical conditions (Gamliel et al., 2020). Simply projecting species distribution models like those derived here under future climate change scenarios is likely to overlook the potential for warming temperatures to exceed thermal optima and limits for species persistence. Moreover, it is critical for studies of vector-borne disease transmission under future climate change to account for the gap between potential and realized future vector thermal niches, as vectors may not immediately track geographic shifts in climate suitability. Approaches that combine mechanistic thermal performance information (e.g., from laboratory studies and life history models) with statistical inference based on current distributions are most likely to accurately capture current and future environmental constraints on species persistence, including for noxious species like disease vectors.

This study incorporates several methodological innovations from previously published species distribution modeling methods for mosquitoes, particularly focused on estimating temperature bounds, with applications for climate change models. First, we computed covariates aligned to the time of collection of our mosquito occurrence data, in contrast to numerous studies that use climatologies estimated from the years preceding the occurrence point sampling time frame (Al Ahmed et al., 2015; Conley et al., 2014; Kraemer et al., 2015; Messina et al., 2019; Mweya et al., 2013; Richman et al., 2018). Second, given our explicit goal of estimating thermal limits, we restricted our measures of average temperature to the relevant activity season of each focal species (Smeraldo et al., 2018), which ensures that we are using the temperature range of the mosquito in the field when that species is active and non-dormant (and therefore comparable to laboratory experimental measurements). Third, we selected background points from the ecoregions in which a species occurs plus a 200 km buffer, as well as adjacent ecoregions, to ensure that we are estimating temperature limitations based on regions that are plausible based on habitat suitability. This method, grounded in ecological theory (Barve et al., 2011; Soberón, 2010; Soberon & Peterson, 2005) and similar to background limitation methods for non-mosquito species in the literature (Gogol-Prokurat, 2011; Khoury et al., 2020; Lira-Noriega et al., 2018; Mertens et al., 2021; Nuñez-Penichet et al., 2019), functionally decouples habitat and temperature. This is largely a new advancement for mosquito species distribution modeling. Finally, while most mosquito species distribution modeling studies focus on identifying predictors of one or two mosquito species, here we explicitly aim to infer thermal limits and optima and to compare these to mechanistic estimates (Brady et al., 2013) among multiple medically-relevant mosquito vector species (Cianci et al., 2015; Cunze et al., 2016; Kraemer et al., 2015; Kulkarni et al., 2010; Uusitalo et al., 2019).

However, our study also has several limitations. First, the lack of thermal breadth in the background sampling may limit our ability to accurately infer thermal maxima.. Mosquito species that occur in hotter climates, such as *Ae. aegypti, An. stephensi,* and *An. gambiae*, lack a sufficient amount of ecoregion-matched background points that have comparably high, or higher, temperatures than the occurrence points, making it difficult to estimate the temperature range that would be hot enough to prevent occurrence (i.e., identify thermal maxima) (Merow et al., 2013). Second, because our goal was to create accurate but comparable species distribution models for seven mosquito species across their global extent, we primarily relied on publicly available GBIF occurrence points (see Methods for data filtering criteria), which may not capture the full extent of each species that other sources such as local vector surveillance data might capture. Third, since we were focused on comparing laboratory-derived thermal performance curves to field occurrence, we used temperatures measured during the activity season as predictors in XGBoost models for four out of seven species. Despite the importance of activity season temperature, other climatic variables such as winter low temperatures or dry-season aridity may be equally or more important limitations on species distributions. Fourth, the species distribution modeling framework assumes that species occupy their full climatic niche, but this may not be the case for species that are actively expanding their ranges. Finally, our analysis is based on a core assumption that using the 2000-2019 average of covariates can accurately characterize the average ecological or habitat preferences of a mosquito reported in a specific date and location.

## CONCLUSIONS

As climate change shifts the geographic and seasonal distribution of environmental conditions, it is critical to understand how temperature limits species ranges. Temperature in particular affects vector-borne diseases because of its effects on vector biting rate, parasite incubation rate, survival, and other life history traits. However, the thermal constraints on vector occurrence are less well understood, and particularly how they vary among important vector species. Here, we showed that temperature mean and variation during the activity season provide important constraints on the ranges of seven important mosquito vectors, and that thermal minima and, to a lesser degree, thermal optima observed in the field are closely correlated to those measured in laboratory experimental studies. This finding suggests that species distribution models can, to some extent, contribute to understanding the thermal biology of organisms that cannot be studied in a laboratory setting. Importantly, statistical species distribution models derived from field observations did not clearly identify upper thermal constraints even though these have been directly demonstrated experimentally in the laboratory. Climate change is likely to drive many regions past the currently observed range of temperatures. This highlights the critical need for mechanistic, trait-based studies that capture temperature ranges at which mosquito abundance and occurrence begin to decline (Mordecai et al., 2013; Ryan et al., 2015, 2019) and, at the very least, emphasizes that extrapolations from statistical models based on current species distributions should be validated using physiological models (Elith & Leathwick, 2009; Hijmans & Graham, 2006; Khatchikian et al., 2011).

## ACKNOWLEDGMENTS

TSA is supported by the National Institute of General Medical Sciences (grant no. T32GM144273). MLC was supported by the Illich-Sadowsky Fellowship through the Stanford Interdisciplinary Graduate Fellowship program at Stanford University and an Environmental Fellowship at the Harvard University Center for the Environment. EAM and CKG were supported by the National Science Foundation and the Fogarty International Center (grant no. DEB-2011147). EAM was additionally supported by the National Institute of Allergy and Infectious Diseases (grant nos R01AI168097 and R01AI102918), the National Institutes of Health (grant no. R35GM133439), and by seed grants from the Stanford Woods Institute for the Environment, King Center on Global Development, Center for Innovation in Global Health and Terman Award. CKG was additionally supported by a Stanford Institute for Human-centered Artificial Intelligence Postdoctoral Fellowship. The authors thank Marta Shocket and Oswaldo Villena for providing mosquito life history trait data, and Marianne Sinka and Joshua Longbottom for supplying additional species occurrence points for *Anopheles stephensi*.

## Notes

### Competing Interest Statement

The authors have declared no competing interest.

### Summary of Updates

Author information has been updated with middle initials and orcid ids.

