## Supplemental materials for "Temperature dependence of mosquitoes: comparing mechanistic and machine learning approaches"

**Table S1. Filtered number of occurrences by species after each data cleaning step.**

| Species | *Ae. aegypti* | *Ae. albo-pictus* | *An. gambiae* | *An. stephensi* | *Cx. pipiens* | *Cx. quinque-fasciatus* | *Cx. tarsalis* |
|---|---|---|---|---|---|---|---|
| **Raw Occurrences** | 37,115 | 39,488 | 13,650 | 1,232 | 98,276 | 30,978 | 44,496 |
| **Known Basis of Record** | 36,981 | 39,449 | 13,650 | 1,196 | 98,248 | 30,964 | 44,487 |
| **Records Within 2000-2019 Range** | 30,956 | 34,399 | 10,116 | 540 | 60,078 | 27,823 | 34,361 |
| **Coordinate Uncertainty Reported** | 30,908 | 34,122 | 10,116 | 540 | 59,661 | 27,744 | 34,358 |
| **Coordinate Uncertainty Decimal Filter** | 30,767 | 34,047 | 10,114 | 540 | 59,661 | 27,738 | 34,358 |
| **Within Landmass** | 30,651 | 33,885 | 10,109 | 535 | 59,629 | 27,640 | 35,358 |
| **Within Activity Season** | 30,651 | 29,750 | 10,054 | 535 | 59,429 | 27,640 | 35,348 |
| **Complete Predictors** | 30,598 | 29,714 | 10,052 | 534 | 59,415 | 27,560 | 31,969 |
| **Unique Cell Centroids** | 9,299 | 8,688 | 903 | 358 | 1,949 | 2,670 | 909 |

**Table S2. Environmental covariates and respective data sources, accessed from Google Earth Engine.**

| Covariate | Acronym | Source | Units | Spatial Resolution †,* | Temporal Resolution ^ |
|---|---|---|---|---|---|
| Cattle Density | CD | Gridded Livestock of the World v3 (Gilbert et al., 2018) | Animals per 1 sq. kilometer | 9.25 km # | Single year (2010) |
| Enhanced Vegetation Index Mean | EVIM | NASA MOD13A2 (Didan, 2015) | EVI | 1 km | 16 days |
| Enhanced Vegetation Index Standard Deviation | EVISD | NASA MOD13A2 (Didan, 2015) | EVI | 1 km | 16 days |
| Forest Cover | FC | NASA MOD44B (Dimiceli et al., 2015) | % of a pixel | 250 m | Yearly |
| Human Population Density | HPD | GHS-POP (Freire et al., 2016) | Persons per 1 sq. kilometer | 1 km | Single year (2000, 2015) |
| Precipitation of the Driest Quarter | PDQ | ERA5 (Hersbach et al., 2020) | Millimeters | 27.83 km | Daily |
| Precipitation of the Wettest Quarter | PWQ | ERA5 (Hersbach et al., 2020) | Millimeters | 27.83 km | Daily |
| Surface Water Seasonality | SW | JRC Global Surface Water (Pekel et al., 2016) | Number of months | 30 m | Single year (2014-15) |
| Temperature Mean in Photoperiod Activity Season | Photo ASTM | ERA5 (Hersbach et al., 2020) | °C | 27.83 km | Daily |
| Temperature Standard Deviation in Photoperiod Activity Season | Photo ASTSD | ERA5 (Hersbach et al., 2020) | °C | 27.83 km | Daily |
| Temperature Mean in Precipitation Activity Season | Precip ASTM | ERA5 (Hersbach et al., 2020) | °C | 27.83 km | Daily |
| Temperature Standard Deviation in Precipitation Activity Season | Precip ASTSD | ERA5 (Hersbach et al., 2020) | °C | 27.83 km | Daily |
| Temperature Mean Year-Round | TAM | ERA5 (Hersbach et al., 2020) | °C | 27.83 km | Daily |
| Temperature Standard | TASD | ERA5 (Hersbach | °C | 27.83 km | Daily |

| | | | | | |
|---|---|---|---|---|---|
| Deviation Year-Round | | et al., 2020) | | | |
| Wind Speed | WS | TerraClimate (Abatzoglou et al., 2018) | Meters per second | 4.64 km | Monthly |

† The following conversions are used to acquire spatial resolution in kilometers:
1 minute of arc ≈ 0.0166667 degrees,
1 degree ≈ 111 kilometers
* All rasters are resampled to 1 km x 1 km grid cells before use as a predictor in the machine learning model
^ All rasters are averaged over the 2000-2019 study period, if adequate temporal data is available
### Sourced as 5 arcmin

**Table S3. Thermal minima, optima, and maxima with rank ordering across species for the mechanistic trait based model (M(T)) versus the species distribution model (partial dependence plots (PDPs)).**

| Species | Model | Thermal Minima (Rank) | Thermal Optima (Rank) | Thermal Maxima (Rank) [#] |
|---|---|---|---|---|
| *Aedes aegypti* | M(T) | 16 °C (2) | 27 °C (1) | 34 °C (3) |
| | PDP | 10.5 °C (3) | 25.6°C (4) | -- |
| *Aedes albopictus* | M(T) | 15.5 °C (3) | 25 °C (2) | 30.5 °C (7) |
| | PDP | 9.2 °C (5) | 23 °C (5) | -- |
| *Anopheles gambiae* | M(T) | 18 °C (1) | 24 °C (3) | 31 °C (5) |
| | PDP | 18.5 °C (1) | 26.5 °C (1) | -- |
| *Anopheles stephensi* | M(T) | 15 °C (4) | 23.5 °C (4) | 37 °C (1) |
| | PDP | 16.5 °C (2) | 26.8 °C (2) | -- |
| *Culex pipiens* | M(T) | 11 °C (6) | 22 °C (6) | 35 °C (2) |
| | PDP | 4.7 °C (6) | 12.5 °C (7) | -- |
| *Culex quinquefasciatus* | M(T) | 13 °C (5) | 22.5 °C (5) | 31 °C (5) |
| | PDP | 11 °C (3) | 22.5°C (2) | -- |
| *Culex tarsalis* | M(T) | 9 °C (7) | 21.5 °C (7) | 34 °C (3) |
| | PDP | 5.9 °C (7) | 11.3 °C (6) | -- |

[#] Thermal maxima were not discernible from the partial dependence plots, and are thus not reported.

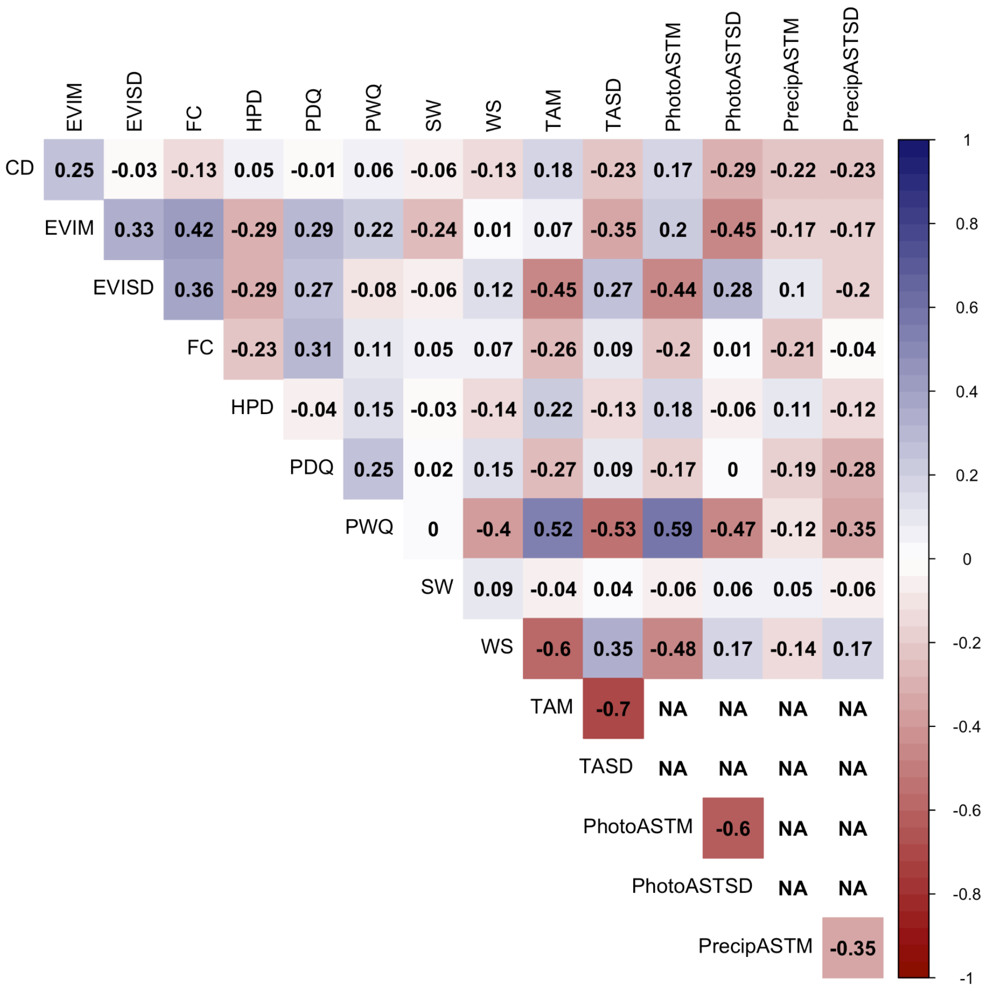


**Figure S1. Pairwise correlation plot of environmental predictors used in the model.** A pairwise correlation analysis was run for each set of two covariates, and any variables that had a correlation that exceeded the R < |0.8| threshold were reassessed and modified, or dropped from the analysis entirely. CD is cattle density; EVIM is enhanced vegetation index mean; EVISD is enhanced vegetation index standard deviation; FC is forest cover percentage; HPD is human population density; PDQ is precipitation of the driest quarter; PhotoASTM is photoperiod activity season temperature annual mean; PhotoASTSD is photoperiod activity season temperature standard deviation; PrecipASTM is precipitation activity season temperature annual mean; PrecipASTSD is precipitation activity season temperature standard deviation; PWQ is precipitation of the wettest quarter; SW is surface water seasonality; TAM is year-round temperature annual mean; TASD is year-round temperature annual standard deviation; and WS is wind speed. There are NAs among temperature variables as there was only one set included in each model (e.g., the *Ae. albopictus* model included PhotoASTM and PhotoASTD but not TAM, TASD, PrecipASTM, or PrecipASTD).

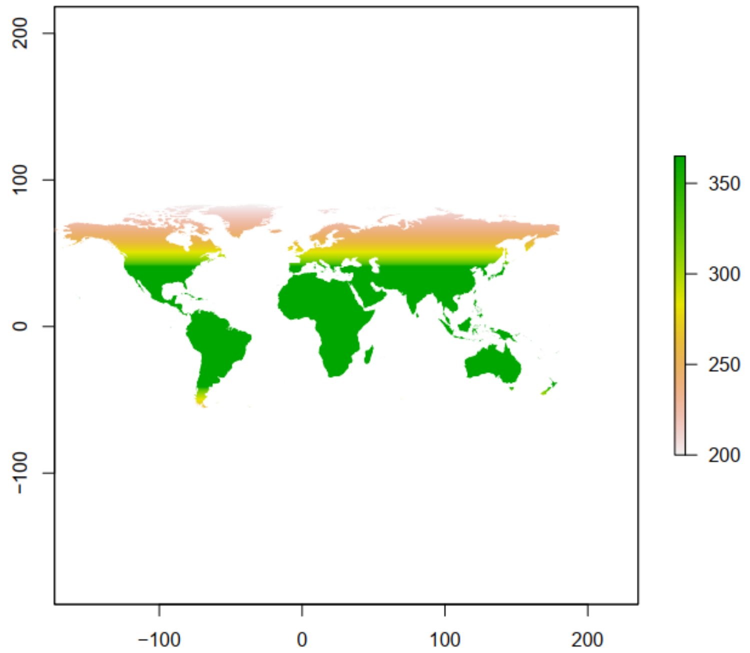

**Figure S2. Photoperiod activity season**. Map of the world shaded in with the length of the photoperiod activity season.

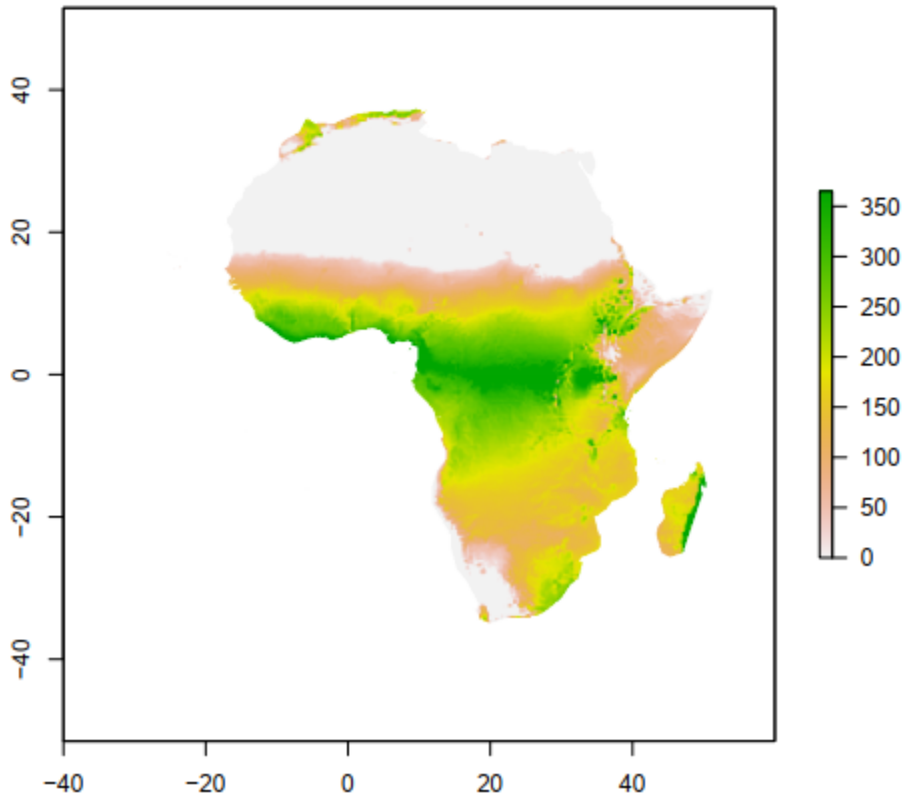


**Figure S3. Precipitation activity season.** Map of Africa shaded in with the length of the precipitation activity season in days.

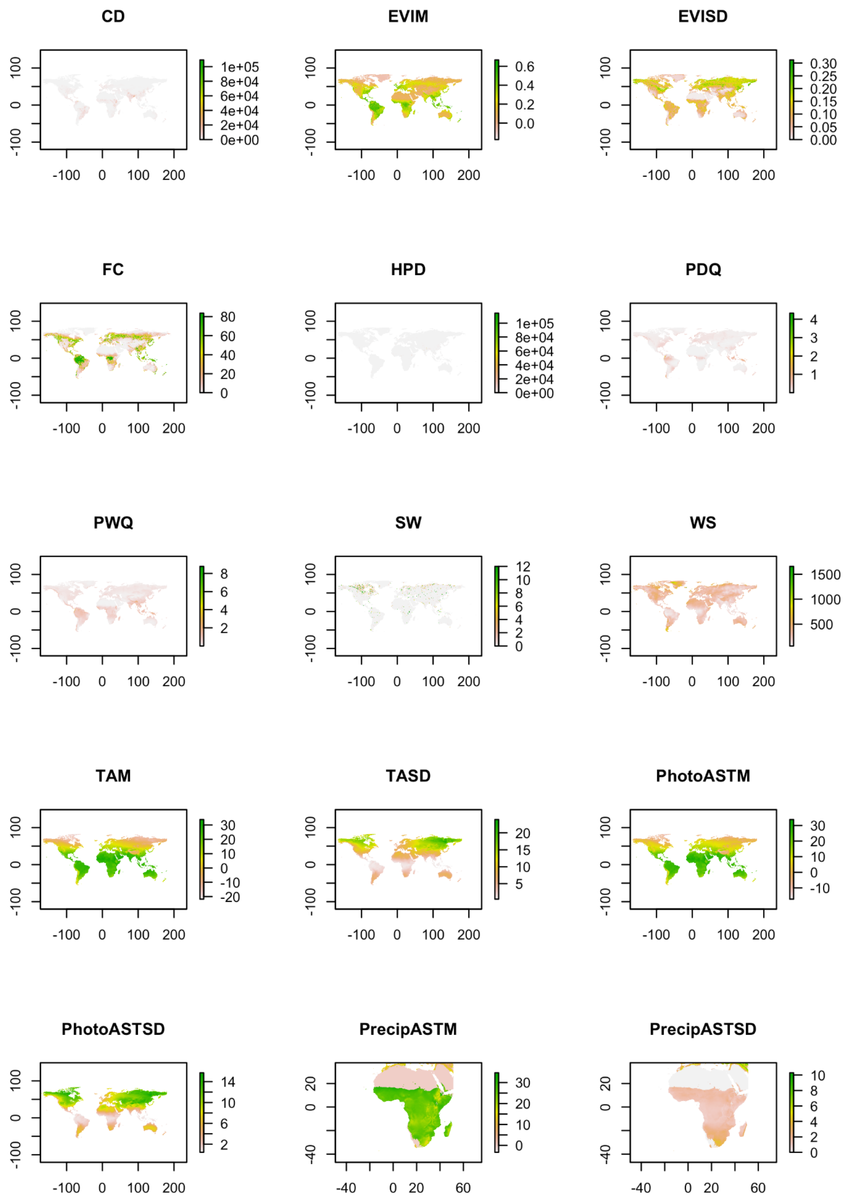


**Figure S4. Raster plots of all environmental covariates used in the model.** Geographic and spatial distribution of every covariate used in the model, including cattle density (CD), enhanced vegetation index mean (EVIM), enhanced vegetation index standard deviation (EVISD), forest cover percentage (FC), human population density (HPD), precipitation of the driest quarter (PDQ), photoperiod activity season temperature mean (PhotoASTM), photoperiod activity season temperature standard deviation (PhotoASTSD), precipitation activity season temperature mean (PrecipASTM), precipitation activity season temperature standard deviation (PrecipASTSD), precipitation of the wettest quarter (PWQ), surface water seasonality (SW), year-round temperature annual mean (TAM), year-round temperature annual standard deviation (TASD), and wind speed (WS). PrecipASTM and PrecipASTSD are only depicted over Africa as this is the region the data was used (*An. gambiae*'s data points are restricted to Africa).

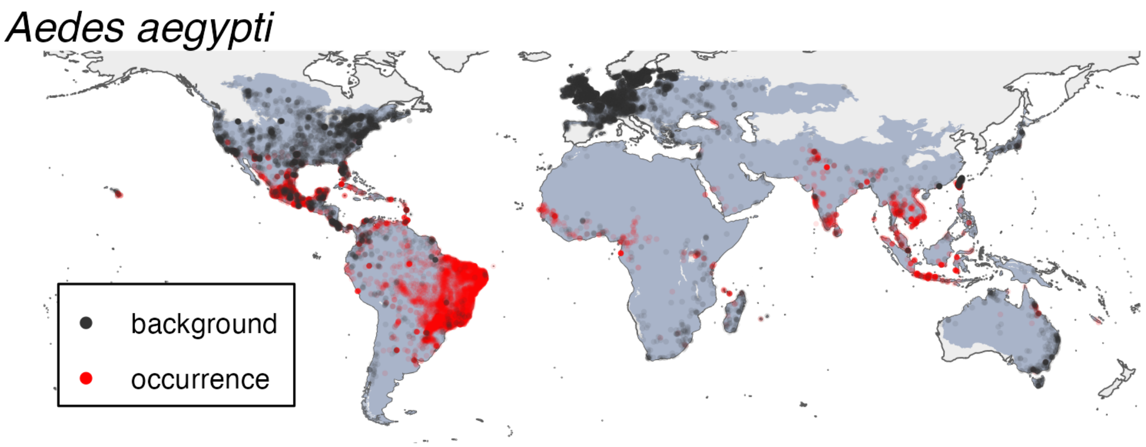


**Figure S5**. **Species occurrence and pseudo-absence background map for *Aedes aegypti*.** Species occurrence centroids (red) and associated pseudo-absence background centroids (black) are plotted, superimposed on the respective set of ecoregions in which their buffered occurrence centroids fall and adjacent ecoregions (gray).

#### *Aedes albopictus*

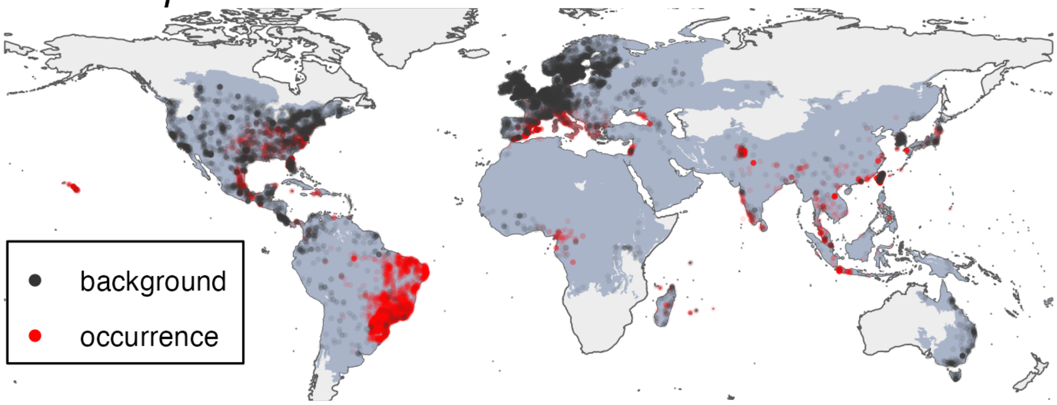


**Figure S6**. **Species occurrence and pseudo-absence background maps for *Aedes albopictus*.** Species occurrence centroids (red) and associated pseudo-absence background centroids (black) are plotted, superimposed on the respective set of ecoregions in which their buffered occurrence centroids fall and adjacent ecoregions (gray).

*Anopheles stephensi*

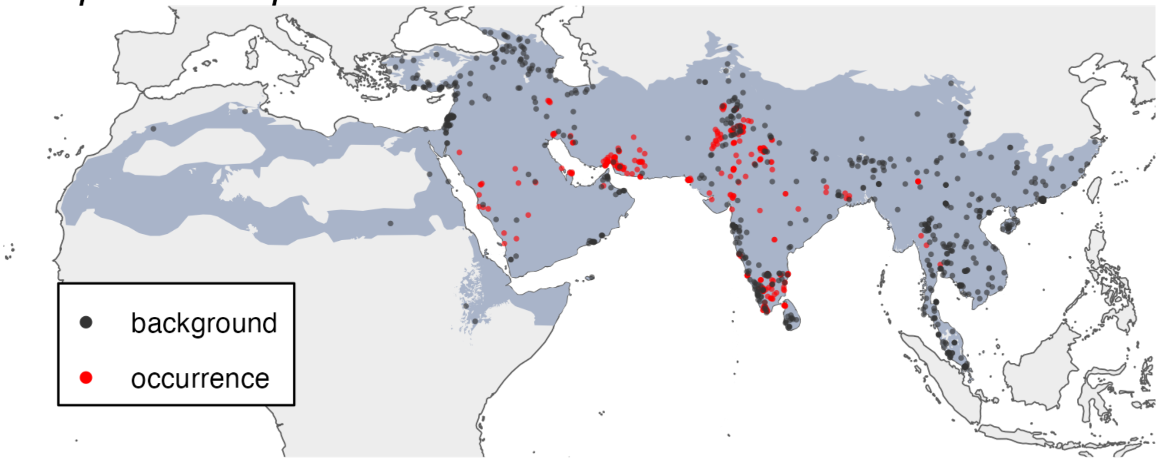


**Figure S7**. **Species occurrence and pseudo-absence background maps for *Anopheles stephensi*.** Species occurrence centroids (red) and associated pseudo-absence background centroids (black) are plotted, superimposed on the respective set of ecoregions in which their buffered occurrence centroids fall and adjacent ecoregions (gray).

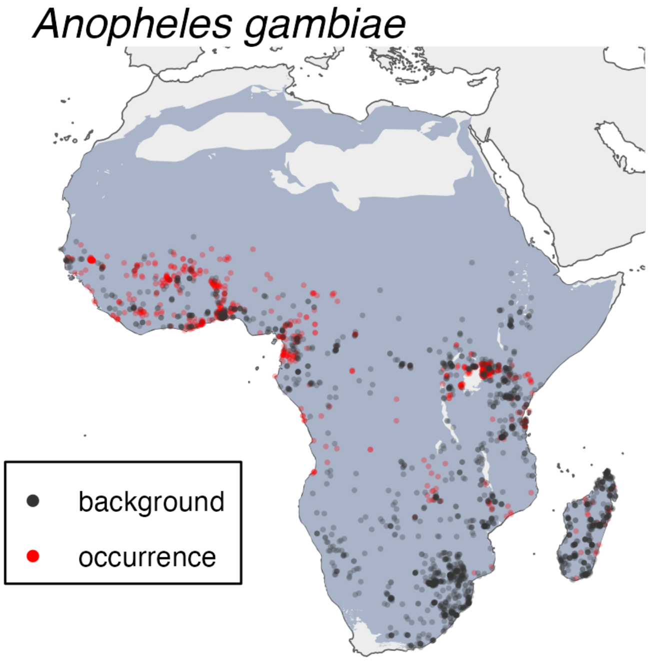


**Figure S8**. **Species occurrence and pseudo-absence background maps for *Anopheles gambiae*.** Species occurrence centroids (red) and associated pseudo-absence background centroids (black) are plotted, superimposed on the respective set of ecoregions in which their buffered occurrence centroids fall and adjacent ecoregions (gray).

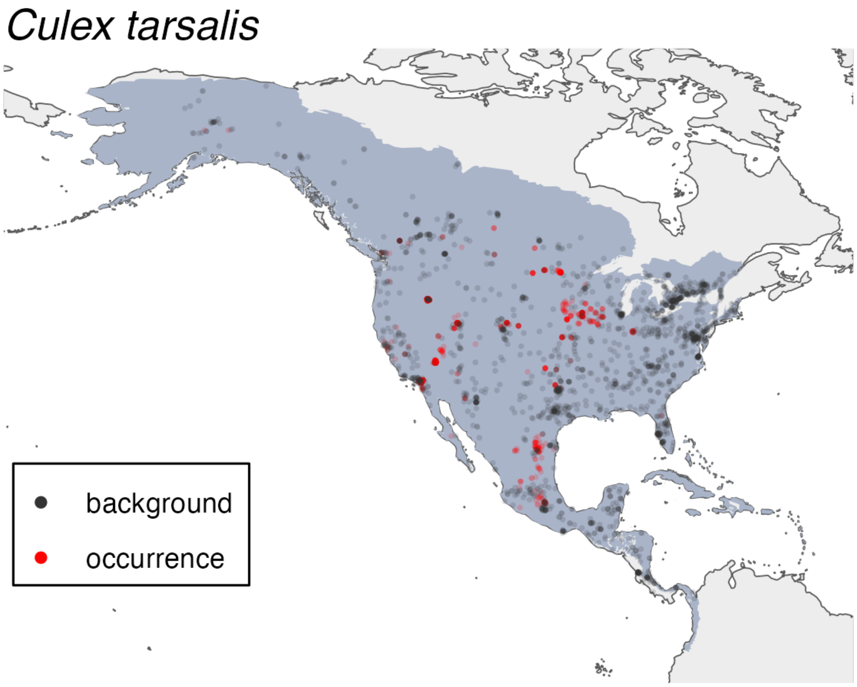


**Figure S9**. **Species occurrence and pseudo-absence background maps for *Culex tarsalis*.** Species occurrence centroids (red) and associated pseudo-absence background centroids (black) are plotted, superimposed on the respective set of ecoregions in which their buffered occurrence centroids fall and adjacent ecoregions (gray).

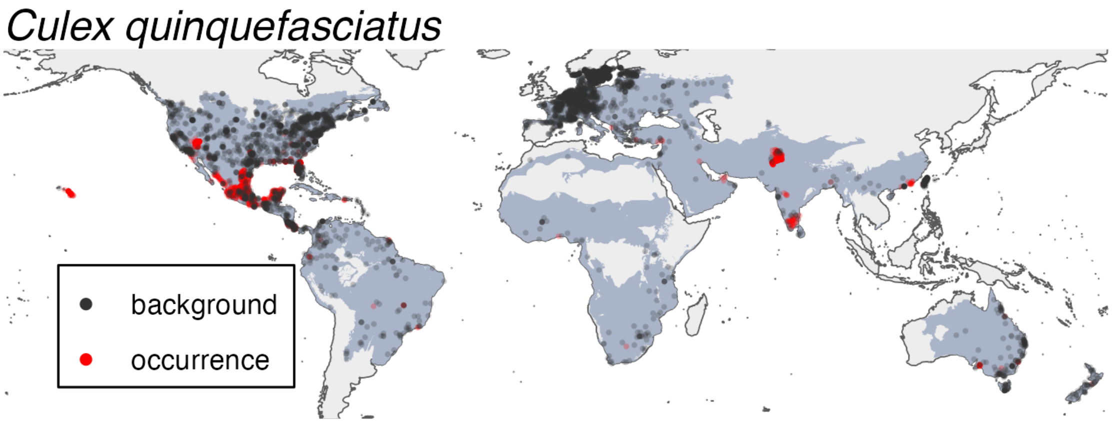


**Figure S10**. **Species occurrence and pseudo-absence background maps for *Culex quinquefasciatus*.** Species occurrence centroids (red) and associated pseudo-absence background centroids (black) are plotted, superimposed on the respective set of ecoregions in which their buffered occurrence centroids fall and adjacent ecoregions (gray).

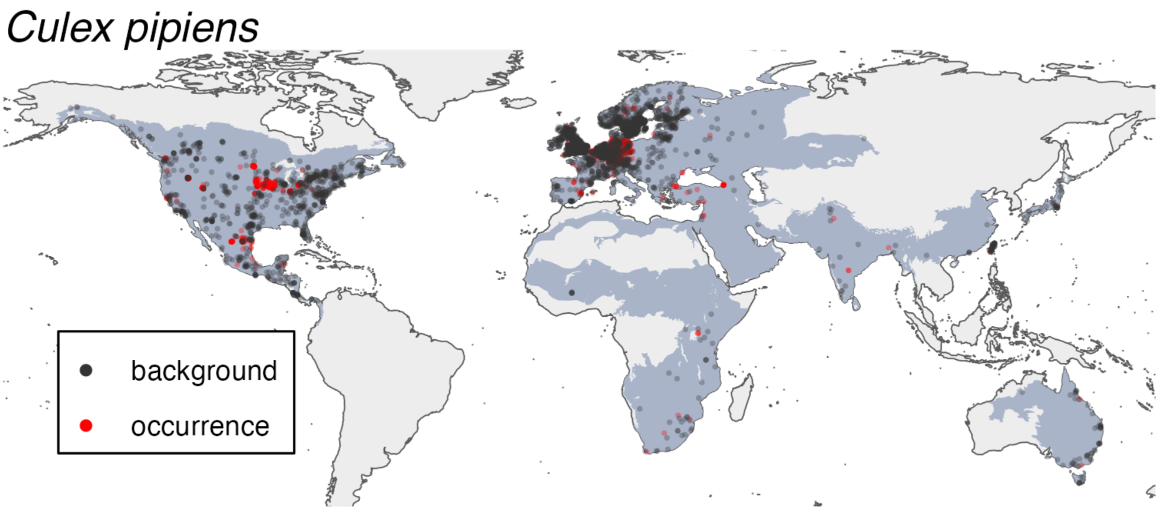


**Figure S11**. **Species occurrence and pseudo-absence background maps for *Culex pipiens*.** Species occurrence centroids (red) and associated pseudo-absence background centroids (black) are plotted, superimposed on the respective set of ecoregions in which their buffered occurrence centroids fall and adjacent ecoregions (gray).

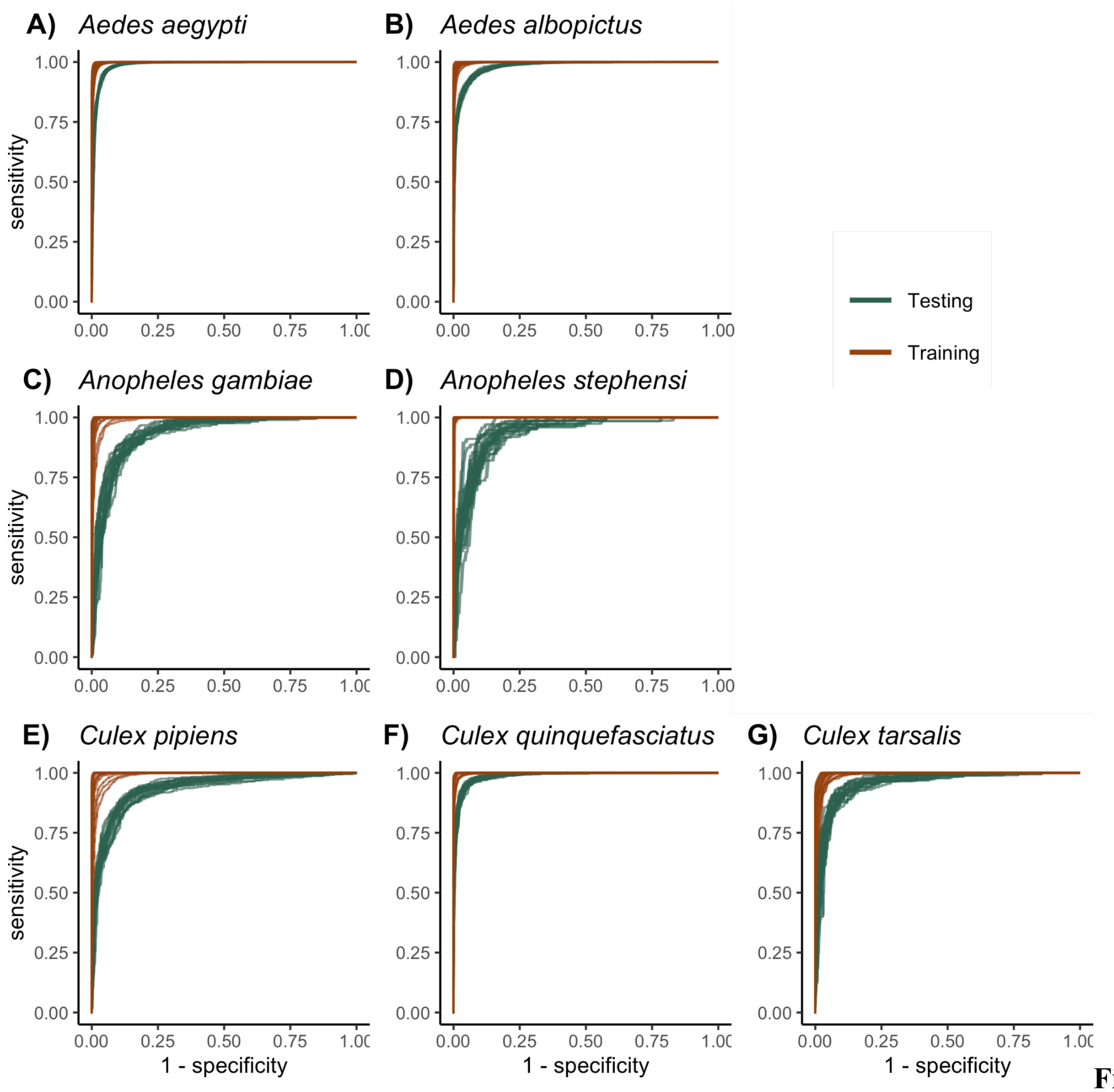


**Figure S12. XGBoost models accurately predicted mosquito occurrence in- and out-of-sample.** Receiver operating characteristic (ROC) curves and area under the curve (AUC) values for assessment of model discrimination are depicted, where 1 represents perfect discrimination of presence and absence and 0.5 represents discrimination no better than chance. Curves depict the training set (red) and the evaluation set (blue).

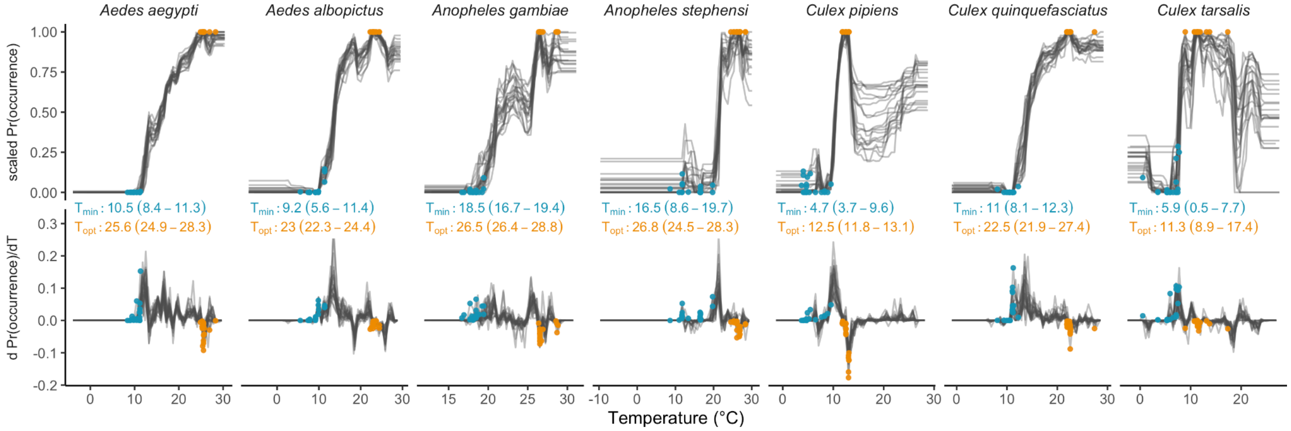


**Figure S13. Thermal minima and thermal optima derived from the PDPs.** Thermal minima were identified as the temperature at which the partial dependence plot began increasing, which we operationalized as the first time the empirical derivative was positive and stayed positive for the next step in temperature as well (Fig. 1c). Thermal optima were identified as the point where the empirical derivative was zero and the partial dependence plot was at its maximum (Fig. 1c). We did not identify thermal maxima, as few species had partial dependence plots that clearly declined after the thermal optima and then reached a lower plateau that was within the range of observed temperatures. Central thermal minima and optima values are medians across the 20 model iterations, and parentheses indicate the full range of values over the model iterations.

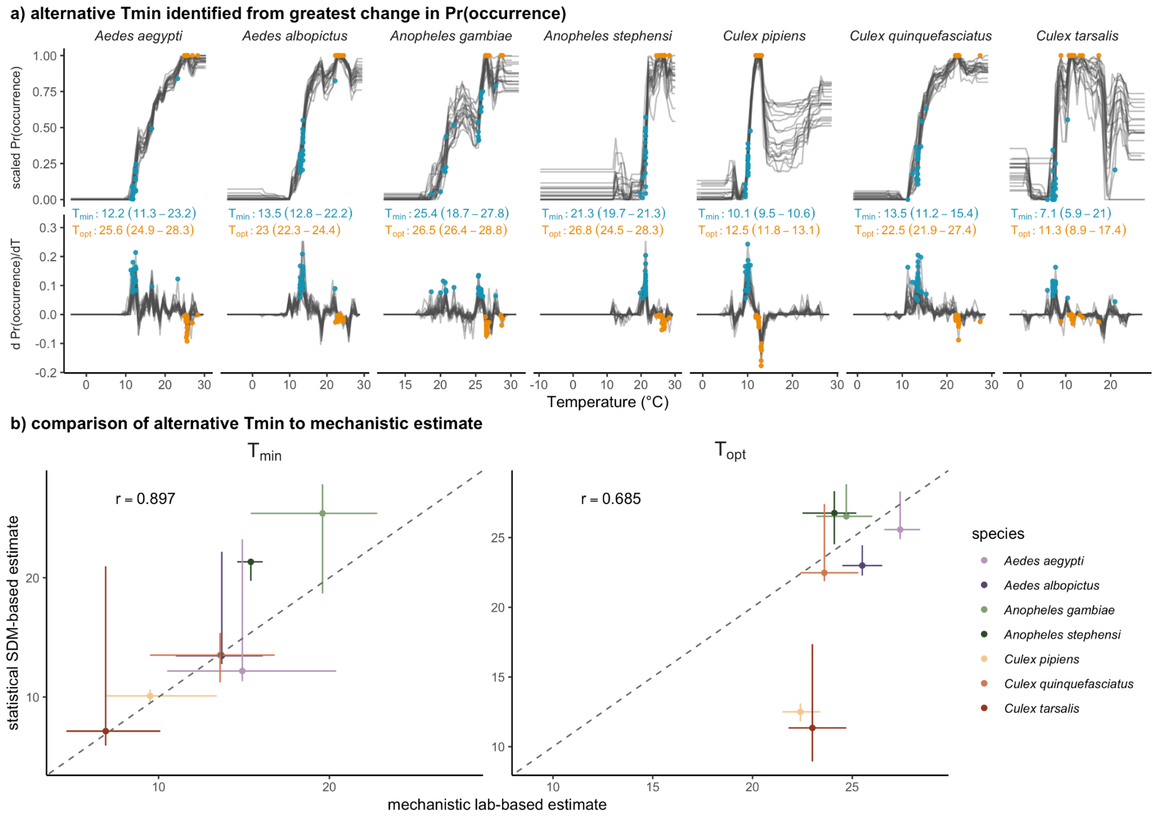


**Figure S14. Comparison between statistical and mechanistic under alternative definition of $T_{min}$.** Lower thermal limits are instead identified as the temperature with the largest empirical derivative in the PDP. Panel a is as in Figure S13. Panel b is as in Figure 4.
